# Mean exercise torque is a critical factor influencing neuromuscular fatigability induced by exhausting contractions

**DOI:** 10.1101/2024.03.06.583664

**Authors:** Loïc Lebesque, Gil Scaglioni, Patrick Manckoundia, Alain Martin

## Abstract

**PURPOSE:** To get a more detailed description of neuromuscular fatigability, maximal torque sustainability (i.e., the ability to maintain a high torque level) can be assessed in addition to the classically used maximal voluntary contraction (MVC). Since this parameter appears to be affected by mean exercise torque (MET), the present study aims to examine the relationship between MET and neuromuscular fatigability induced by exhausting contractions.

**METHODS:** Thirteen participants sustained a plantar flexors MVC for 1 min (MVC_1-MIN_) before and after exhausting exercises designed to produce a similar MET (30% MVC), and following a 10-min rest period. Exercises consisted of intermittent (INT), continuous (CON) or variable (continuous contraction alternating between moderate and low intensity, VAR) contractions performed until task failure.

**RESULTS:** Although the INT resulted in greater exercise duration and torque-time integral than CON and VAR, MVC similarly decreased after all exercises due to neural and muscular impairments. The torque loss during the MVC_1-MIN_ increased after all exercises to a similar extent, mainly because of neural alterations. Contrary to MVC, the torque loss during the MVC_1-MIN_ returned to baseline value after the recovery period.

**CONCLUSION:** By considering both maximal torque production and sustainability, INT, CON and VAR exercises, performed with identical mean torque and until exhaustion, led to a similar neuromuscular fatigability. Results confirm the independence of maximal torque production from the contraction pattern and support the impact of MET on maximal torque sustainability. The present findings are crucial to consider for the management of neuromuscular fatigability in both athletes and patients.

## INTRODUCTION

Neuromuscular fatigability corresponds to the decrement in magnitude or rate of change in a performance criterion, relative to a reference value, over a given time of task performance (Skau et al. 2021). The reduction of the capacity to produce the maximal torque during a brief (i.e., a few seconds long) maximal voluntary contraction (MVC) is the most commonly used index to quantify neuromuscular fatigability. Indeed, a substantial body of studies used the decrease in MVC to investigate the neuromuscular fatigability following a continuous (CON) contraction performed until task failure (TF) with various task modalities, such as different exercise feedback types (Hunter et al. 2002, 2005, 2008; Place et al. 2006a, b, 2007; Klass et al. 2008; Baudry et al. 2009, 2011; Rudroff et al. 2010; Williams et al. 2014), muscle lengths (Place et al. 2005), muscle groups (Neyroud et al. 2013), or even kinds of limb support (Yoon et al. 2009). Although the exercise duration (ED) and torque-time integral (TTI) of the CON exercises seem to be modulated by such exercise parameters, the reduction in MVC was of a similar extent in all studies. The present observation suggests that neuromuscular fatigability, quantified by the decrease in maximal torque production, was similar among task modalities when exercises were performed to exhaustion (i.e., until TF). However, little is known about the influence of contraction patterns on the neuromuscular fatigability induced during such exercises. To address this issue, one of our recent studies compared the neuromuscular fatigability induced by CON and intermittent (15 s on, 5 s off; INT) contractions of plantar flexors (PF) performed at 40% MVC until TF, using a 1-min sustained MVC (MVC_1-MIN_) rather than a brief MVC (Lebesque et al. 2023). As a matter of fact, the MVC_1-MIN_ allows, at the same time, the assessment of two distinct and complementary characteristics of neuromuscular fatigability – the maximal torque production and sustainability (i.e., the ability to sustain the maximal torque) – thus it has the advantage of providing a more detailed and comprehensive description of neuromuscular fatigability (Lebesque et al. 2022, 2023). Although at exhaustion, the ED of the INT exercise was longer than that of the CON, both exercises led to a similar MVC decline, while the maximal torque sustainability was affected by the CON. Based on these results, a CON exercise appears to be more fatiguing than intermittent contractions, when performed at the same intensity until exhaustion. In addition, it is worth noting that the change in maximal torque sustainability was positively correlated to the mean exercise torque (MET). In other words, the greater the mean torque developed during the exhausting exercise, the more the ability to sustain the maximal torque is impaired. This finding suggests that exercises developing an identical MET should lead to similar neuromuscular fatigability.

Combined with MVC, maximal torque sustainability provides a more accurate assessment of the fatiguing nature of exercises performed until exhaustion. It is therefore crucial to acquire information on this neuromuscular capacity and in particular on its relationship with the MET. Thus, the current study aims to further analyse the impact of MET on maximal torque sustainability by extending this analysis to additional exercise profiles generating a similar MET. Therefore, we used CON, INT and variable (VAR) contractions with a MET set at 30% MVC and performed until TF. Considering exhaustion achieved at the end of each exercise, it is hypothesized that the decline in MVC is similar for the three contraction patterns. Since the MET is identical for all exercises, we also expect maximal torque sustainability to be equally compromised by each of them. Finally, because neuromuscular fatigability is a transient phenomenon, the evolution of both capacities was also explored after 10 min of rest. We hypothesize that 10 min of rest after reaching exhaustion is sufficient for complete recovery in maximal torque sustainability but not in maximal torque production, as already observed 10 min after CON and INT contraction at 40% MVC until TF (Lebesque et al. 2023).

## METHODS

### Participants and ethics

Thirteen healthy male volunteers (age = 25 ± 3 years old; body mass = 75.4 ± 8.5 kg; height = 179.0 ± 5.8 cm, body mass index = 23.5 ± 1.8 kg·m^-2^) were enrolled in this study after giving their oral and written informed consent. Participants were recreationally active and none of them suffered from neurological or physical disorders. For all participants, the dominant leg (i.e., the leg used to kick a ball on a target; van Melick et al. 2017) was the right one. This study was carried out in accordance with the latest revision of the Declaration of Helsinki. The CPP SOOM III ethics committee approved the protocol (number 2017-A00064-49).

### Protocol

#### Experimental design

Participants took part in three experimental sessions randomly administered. Sessions lasted approximately 1.5 hours. For each subject, sessions were interspaced by a 7-day interval and planned at the same time of the day to avoid any circadian bias (Guette et al. 2005; Tamm et al. 2009; Douglas et al. 2021). Each session included neuromuscular assessments (torque and electromyographic (EMG) recordings) of PF before (Pre) and immediately after (Post) a submaximal fatiguing exercise, as well as after a 10-min recovery period (Post-10). The fatiguing exercise started 10 min after the completion of the pre-exercise neuromuscular evaluation and it consisted of CON, INT or VAR contractions with a mean torque of 30% MVC performed until exhaustion. Figure 1A shows the experimental protocol used in the present study.

**Figure 1.**
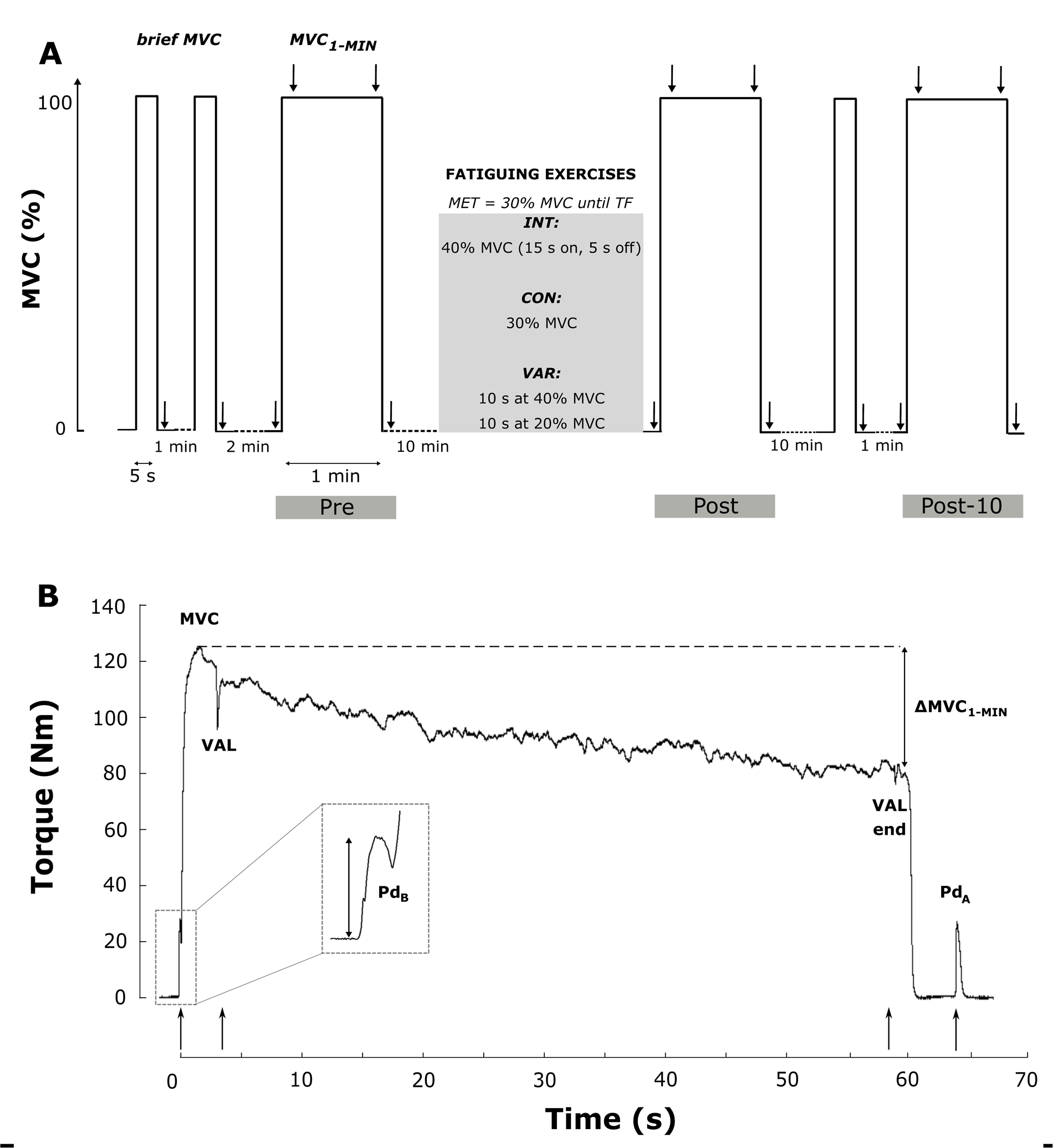
Sum up of the experimental design of the study. A. The protocol consisted in a MVC_1-MIN_ performed before (Pre), after (Post) the fatiguing exercise and after a 10-min recovery period (Post-10). The fatiguing exercise consisted of INT, CON or VAR contractions performed with an equivalent MET (30% MVC) until TF. Note that brief MVCs were performed only at Pre and Post-10. Arrows represent electrical doublets (100 Hz) delivered during the protocol. B. Experimental data of torque during the MVC_1-MIN_ from one typical subject. Main neuromuscular measurements are reported (MVC and ΔMVC_1-MIN_, VAL and VAL end, Pd_B_ and Pd_A_). Note that Pd_B_ was obtained from doublet delivered before the MVC_1-MIN_, only at Post. Pd_B_ at Pre and Post-10 were obtained after the brief MVC. Arrows represent electrical doublets (100 Hz) delivered at rest and superimposed to the contraction. CON = continuous exercise; INT = intermittent exercise; MET = mean exercise torque; MVC = maximal voluntary contraction; MVC_1-MIN_ = 1-min sustained MVC; Pd_A_ = potentiated doublet after the MVC_1-MIN_; Pd_B_ = potentiated doublet before the MVC_1-MIN_; TF = task failure; VAL = voluntary activation level; VAR = variable exercise; ΔMVC_1-MIN_ = torque loss during the MVC_1-MIN_

#### Neuromuscular assessment

A standardized warm-up protocol for PF preceded the neuromuscular assessment and consisted of 10-15 PF isometric contractions lasting 5 s each, with an incremental intensity from 25 to 100% of their maximal effort. After warming up, subjects performed two brief PF MVCs (5-s long, with 1-min rest in-between) and 2 s after each contraction an electrical doublet was delivered to obtain the potentiated doublet (Pd). If the difference between the MVCs was > 5%, an additional maximal contraction was performed. Then, 2 min of rest was given to the subjects before the Pre-exercise MVC_1-MIN_. For each MVC_1-MIN_, an electrical doublet was delivered at rest and gave subjects, inter alia, the signal to start the MVC_1-MIN_. Electrical doublets superimposed to the sustained maximal contraction were delivered 3 and 58 s after the MVC_1-MIN_ start. A final doublet was delivered at rest, 3 s after the end of the MVC_1-MIN_. Participants were informed of the duration of the sustained MVC and instructed to perform a maximal performance during the entire contraction in order to lessen the adoption of a pacing strategy (Halperin et al. 2014) during the MVC_1-MIN_. In addition, strong and loud verbal encouragements were given to the subjects to ensure maximal performance during the sustained contraction. The MVC_1-MIN_ were performed before (Pre) and after (Post) the fatiguing exercise and after the 10-min recovery period (Post-10), while the brief MVC was carried out only at Pre and Post-10 because the neuromuscular fatigability should be assessed as soon as possible after the exercise completion (Mira et al. 2017; Place and Millet 2019).

#### Electrical stimulation

Electrophysiological and torque responses of the triceps surae muscles were evoked by percutaneously stimulating the posterior tibial nerve, in the popliteal fossa of the dominant leg. Monophasic rectangular pulses (1 ms) were delivered using a Digitimer stimulator (DS7AH, Digitimer, Hertfordshire, UK), triggered by a commercially available software (Tida, Heka Elektronik, Lambrecht/Pfalz, Germany). The anode (5 x 10 cm, Compex SA, Ecublens, Switzerland) was placed beneath the patella. A hand-held cathode ball (5 mm diameter) was used to locate the optimal stimulation site, namely the site that evoked the greatest soleus (SOL) M-wave amplitude. Once the stimulation site was determined, the cathode (a self-adhesive electrode with a 7 mm diameter, Contrôle Graphique Medical, Brie-Compte-Robert, France) was fixed to this site with tape. A hand-made hard-pad (a cotton cube wrapped in adhesive tape; 4 x 2 x 2 cm) was then positioned over the cathode and firmly fixed with adhesive tape to the limb to keep constant pressure on the electrode throughout the experiment. To determine the intensity at which the M-wave amplitude plateaued (i.e., no further increase in the M-wave amplitude was evident in SOL), the intensity of stimulation was gradually increased by 5 mA from the motor threshold to the maximal M-wave of SOL. This intensity was further increased by 20% to ensure supramaximal stimulations during the entire session and was kept constant throughout the experimental session (mean supramaximal intensity: 72.2 ± 22.0 mA, range: 42 – 144 mA). Only electrical doublets at 100 Hz (1 ms pulse duration, supramaximal intensity) were delivered during the experiment.

#### Fatiguing exercises

Visual feedback of the torque exerted by the subjects and the target torque were displayed on a computer screen placed 1 m in front of them. Fatiguing exercises consisted in isometric contractions of PF. For the CON session, subjects were asked to sustain the target torque (30% MVC) until TF. For the INT session, subjects were instructed to perform intermittent (15 s on, 5 s off with sound signals; duty cycle: 75%) contractions at 40% MVC until TF. Finally, for the VAR session, subjects were required to perform a sustained contraction characterized by alternating periods of 10 s at 20% MVC and 10 s at 40% MVC (i.e., variable contraction intensity) until TF. It is noteworthy that, for this contraction pattern, half of the subjects started the exercise with 10 s at 20% MVC while the other half started at 40% MVC. In each session, the expected MET was 30% MVC. The TF was defined as a drop of the torque below the target for three consecutive seconds. All subjects were strongly encouraged throughout the entire exercise and naive about the criterion leading to exercise interruption. The rate of perceived exertion (RPE), defined as “the conscious sensation of how hard, heavy, and strenuous a physical task is” (Marcora 2010; Pageaux 2016) was assessed at TF for all exercises. Experimental sessions were administered in a randomized order.

### Data acquisition

#### Torque recordings

PF isometric torque was recorded using an isokinetic dynamometer (Biodex System 4 pro®, Biodex Medical System Inc., Shirley, NY, USA). Subjects were comfortably seated on the dynamometer chair with the dominant foot securely strapped to the footplate at the ankle level. The centre of rotation of the dynamometer shaft was aligned with the lateral malleolus. The ankle, knee, and hip angles were set at 90°, 120°, and 130° respectively (180° full extension). To limit trunk movements that could affect torque development, the trunk and hip were firmly strapped to the seat with belts. The setting configuration was kept constant throughout the entire experimental protocol and among sessions. Torque data were recorded at a sampling frequency of 2 kHz with the Biopac acquisition system (Biopac MP160, Biopac System Inc, USA).

#### Electromyographic recordings

The EMG activity was recorded from four muscles of the dominant leg: the medial (MG) and lateral (LG) gastrocnemii, the soleus (SOL), and the tibialis anterior (TA). The EMG signals were recorded bipolarly using silver chloride circular surface electrodes (7 mm recording diameter; Contrôle Graphique Medical, Brie-Compte-Robert, France) with an inter-electrode centre-to-centre distance of 20 mm. The skin was shaved, abraded, and cleaned with alcohol before electrode placement to minimize skin impedance (< 5 kΩ). For SOL, electrodes were placed 3 cm below the insertion of the two gastrocnemii over the Achilles tendon. For MG and LG muscles, electrodes were positioned over the muscle belly. For TA, they were placed on the upper third of the distance between the fibula head and the tip of the lateral malleolus. The reference electrode was positioned between the two gastrocnemii bellies of the same leg in a central position. EMG signals were amplified (gain = 1000) and filtered (10 – 500 Hz). EMG data were recorded at a sampling frequency of 2 kHz with the Biopac acquisition system (Biopac MP160, Biopac System Inc, USA; RRID: SCR_014829).

### Data analysis

Torque and electrophysiological data were processed using custom programs written in Matlab (version 9.4 R2018a, Mathworks, Natick, MA, USA; RRID: SCR_001622). An experimental torque trace of the main neuromuscular parameters from a representative subject is depicted in Figure 1B. Data processing was similar to our previous study (Lebesque et al. 2023).

#### Torque analysis

The maximal torque developed in the first 5 s of the MVC_1-MIN_ is a reproducible and reliable index of maximal torque production (Lebesque et al. 2022, 2023) and was then labelled ‘MVC’. The Pre-exercise MVC was used to set the target torque of the fatiguing exercises. The torque loss during each MVC_1-MIN_ (ΔMVC_1-MIN_) was calculated as the relative change between the MVC and the mean torque during the final 2 s of contraction:

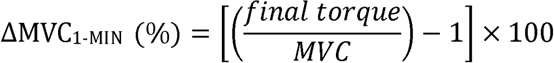

This index of maximal torque sustainability showed good intra- and inter-individual reproducibility (Lebesque et al. 2023). Change in ΔMVC_1-MIN_ between pre and post-exercise was quantified as the difference between both measures:

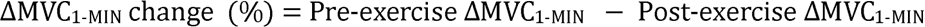

Concerning the potentiated doublets, we measured their amplitude before (Pd_B_) and after (Pd_A_) the MVC_1-MIN_. The Pd_B_, at Pre and Post-10, corresponds to the potentiated doublet elicited just after the brief MVC, while at Post it corresponds to the potentiated doublet elicited just before the MVC_1-MIN_. This latter was potentiated by the fatiguing exercise performed until exhaustion. At each time point (Pre, Post and Post-10), the ratio between the Pd_A_ and the Pd_B_ represents the evolution of the potentiated doublet following each MVC_1-MIN_ (ΔPd_1-MIN_), and was expressed as a percentage of change as follows:

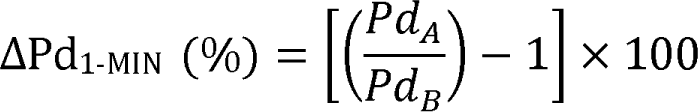

According to the original formula from Allen et al. (1995), the voluntary activation level (VAL) was determined at the beginning (i.e., after 3 s of contraction) and the end (i.e., after 58 s of contraction) of the MVC_1-MIN_. Strojnik and Komi correction (Strojnik and Komi 1998) was applied when the superimposed doublet was not exactly delivered over the peak torque as follows:

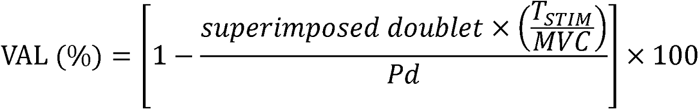

T_STIM_ represents the voluntary torque averaged over a 500 ms time window prior to the superimposed doublet stimulation. Note that the VAL at the end of the MVC_1-MIN_ was corrected using the mean torque of the last 2 s of contraction instead of the MVC in the VAL equation. A value of 100% indicates a full voluntary activation. To calculate the VAL at the beginning of the MVC_1-MIN_, we used the amplitude of the Pd_B_, while at the end of the MVC_1-MIN_, we used the amplitude of the Pd_A_. The ratio between the VAL at the beginning and the end of the MVC_1-MIN_ represents the evolution of the VAL during the MVC_1-MIN_ (ΔVAL_1-MIN_) and was expressed in percentage of change as follows:

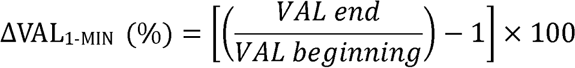

The TTI of the fatiguing exercise, namely the effective amount of torque produced by PF, is computed as the integral of the torque during the entire exercise period minus the integral of the baseline torque at rest during the same period. INT exercise period started with the raising phase of the first contraction and ended after the decline phase of the last contraction. The total ED refers to the duration of the entire exercise (contractions and rest periods). The effective ED represents the real duration of contraction (i.e., without rest periods). Thus, the effective ED is equal to the total ED for CON and VAR exercises. The MET, calculated as the exercise TTI divided by the total ED, corresponds to the mean torque developed during the entire ED.

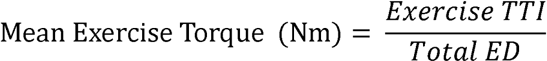

*EMG analysis*. We analysed the second M-wave of each Pd (Kawakami et al. 2000; Matkowski et al. 2011). The peak-to-peak amplitude of the maximal M-wave of the SOL, MG and LG was measured and then summed to determine a comprehensive PF maximal M-wave (M_MAX_). The EMG root mean square (RMS) of the SOL, MG and LG was calculated and summed at both the beginning and the end of the fatiguing exercise. For each measure, RMS was averaged over a 5 s time window. For CON, we consider the RMS between the 5^th^ and the 10^th^ s of the exercise, and between 10 and 5 s before the exercise end. For INT, we consider the RMS between the 5^th^ and the 10^th^ s of the first and last contraction. For VAR, we consider the RMS between 2.5 and 7.5 s of the first and the last contraction of each intensity (i.e., 20 and 40% MVC). For the latter exercise, we averaged the RMS obtained at 20 and 40% MVC at both the beginning and the end of the exercise. Then, the RMS, at the beginning and the end of each fatiguing task, was normalized by the Pre and Post M_MAX_ (RMS/M_MAX_) respectively. The ratio between the RMS/M_MAX_ at the beginning and the end of the exercise represents the evolution of the muscle activities during the fatiguing task and was expressed in percentage of change as follows:

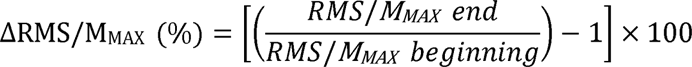

### Psychometric analysis

The RPE was quantified at TF using the CR-100 Scale (Borg 2007).

### Statistical analysis

It was estimated that thirteen participants were needed to detect differences with moderate effect size (η_p_² = 0.08) using standard parameters of 1-β = 0.95 and α = 0.05. We used the Shapiro-Wilk test to assess the normality of each dataset. Because all variables follow the normal distribution, we used only parametric analyses of variance (ANOVA). Factors used in ANOVAs were *pattern* (INT, CON and VAR), *time* (Pre, Post and Post-10) and *duration* (beginning *vs*. end of fatiguing exercise). One-way ANOVAs (*pattern* factor) were performed on total and effective ED, exercise TTI, MET, RPE at TF and ΔRMS/M_MAX_. Two-way repeated-measures ANOVAs (*pattern* and *time* factors) were conducted on MVC, VAL, M_MAX_, Pd_B_, ΔMVC_1-MIN_, ΔVAL_1-MIN_ and ΔPd_1-MIN_. A two-way repeated-measures ANOVA (*pattern* and *duration* factors) was performed on RMS/M_MAX._ When a significant effect was observed, we used Tukey’s test for post-hoc comparisons. When necessary, we performed Bayesian equivalence analysis with a region of practical equivalence ROPE = [-0.1:0.1] and a prior Cauchy scale of 0.707 (Morey and Rouder 2011). The correlation between muscle activity during the fatiguing exercise (RMS/M_MAX_) and the effective ED was analysed using repeated-measures correlation (rmcorr R package, Bakdash and Marusich 2017). Effect sizes of ANOVAs are reported as partial eta squared (η_p_²) with small (≥ 0.01), moderate (≥ 0.07) and large effects (≥ 0.14). Results are reported as mean ± standard deviation. We set the significant level at *p* < 0.050. The sample size was estimated using G*Power software (version 3.1.9.2, Universität Düsseldorf, Germany; RRID: SCR_013726). The repeated-measures correlation was performed and plotted with R (R Core Team, 2017, version 4.1.3; RRID: SCR_001905). Bayesian equivalence tests were conducted with JASP software (JASP Team, 2023, version 0.16; RRID: SCR_015823). We conducted all inferential statistical analyses using Statistica software (Statsoft, version 13, Tulsa, OK, USA; RRID: SCR_014213). Graphs were made with GraphPad Prism software (GraphPad Software Inc., version 8.4.0, San Diego, CA, USA; RRID: SCR_002798).

## RESULTS

### Parameters characterizing fatiguing exercise

The MET, a-priori set at 30% MVC, was not different among exercises, whether expressed in absolute (F_(2,24)_ = 0.600, *p* = 0.557, η_p_² = 0.048) or relative (F_(2,24)_ = 0.720, *p* = 0.496, η_p_² = 0.057) values (Table 1). Bayesian equivalence tests, among exercises MET, showed that the overlapping hypothesis Bayes Factor (BF^OH^_01_) was superior to 2.137 and the non-overlapping hypothesis Bayes Factor (BF^NOH^_01_) was superior to 2.406, meaning that the data are 2.1 times more likely to validate the null hypothesis than the alternative one and 2.4 times more likely to lie in the equivalence than in the non-equivalence region. For the INT exercise, participants performed 52.5 ± 27.2 contractions. The analysis of variance detected a significant effect of *pattern* on total ED (F_(2,24)_ = 23.686, *p* < 0.001, η_p_² = 0.664) and effective ED (F_(2,24)_ = 12.070, *p* = 0.002, η_p_² = 0.501). The INT total and effective ED were longer than CON (+106.0 %, *p* < 0.001 and +54.7%, *p* < 0.001, respectively) and VAR (+93.7 %, *p* < 0.001 and +45.5%, *p* = 0.002, respectively) (Table 1). No difference in ED between CON and VAR was spotted (*p* = 0.862; Bayesian equivalence tests: BF^OH^_01_ = 3.349, BF^NOH^_01_ = 4.354) (Table 1). We found a main *pattern* effect on exercise TTI (F_(2,24)_ = 42.404, *p* < 0.001, η_p_² = 0.779). The TTI of the INT exercise was greater compared to that of CON (+104.9%, *p* < 0.001) and VAR (+86.8%, *p* < 0.001) contractions (Table 1), with no difference between CON and VAR values (*p* = 0.725). We found a significant *pattern* x *duration* interaction effect (F_(2,24)_ = 4.471, *p* = 0.022, η_p_² = 0.271) on RMS/M_MAX_. At the beginning of the fatiguing exercise, the RMS/M_MAX_ of the triceps surae muscles was greater for INT compared to CON (*p* = 0.002) and VAR (*p* = 0.017) while at the end of the fatiguing exercise, no more difference was observed among patterns (*p*-range: 0.871 – 1.000). Compared to the initial value, this parameter significantly increased at the end of each fatiguing exercise (INT: +44.0%, *p* < 0.001; CON: +120.7%, *p* < 0.001; VAR: +87.2%, *p* < 0.001). In addition, we found a significant negative correlation between ΔRMS/M_MAX_ and the effective ED (r*_rm_*(25) = -0.391, *p* = 0.044, CI_95%_ = [-0.671 : 0.013]) (Figure 2). Experimental torque and EMG traces during exercises, for a representative subject, are depicted in Figure 3.

**Figure 2.**
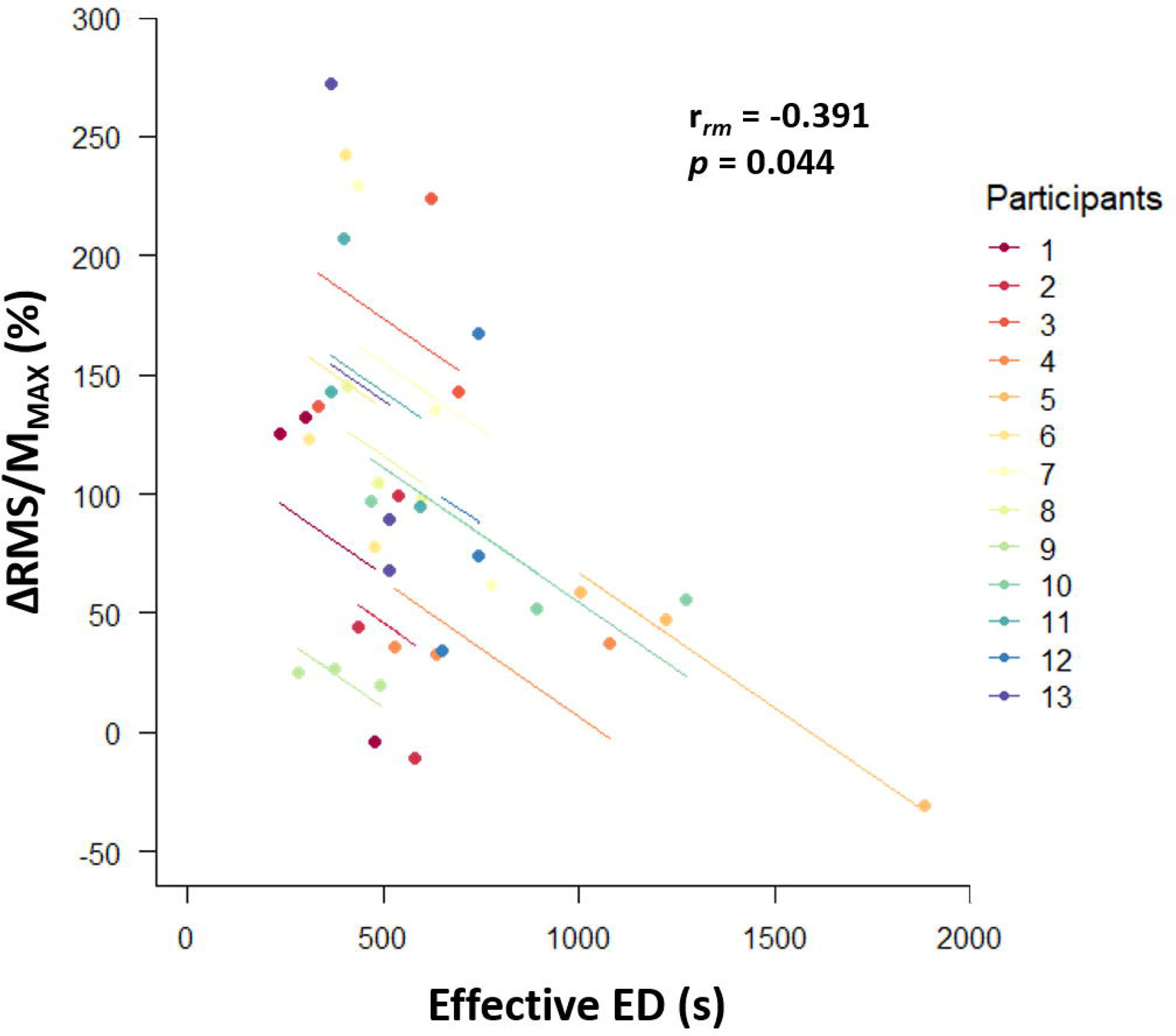
Individual repeated-measures correlation between effective ED and change in electromyographic activities during exercise. The lines correspond to the repeated- measures correlation fit for each participant (*n* = 13). For each fatiguing exercise, data from the same participant are shown in the same colour. ΔRMS/M_MAX_ represents the change in muscle activities, normalized by M_MAX_, between the beginning and the end of the fatiguing exercise. ED = exercise duration; M_MAX_ = sum of maximal M-wave of triceps surae muscles; RMS = sum of electromyographic root mean square of triceps surae muscles. *p* < 0.050 expresses a significant correlation

**Figure 3.**
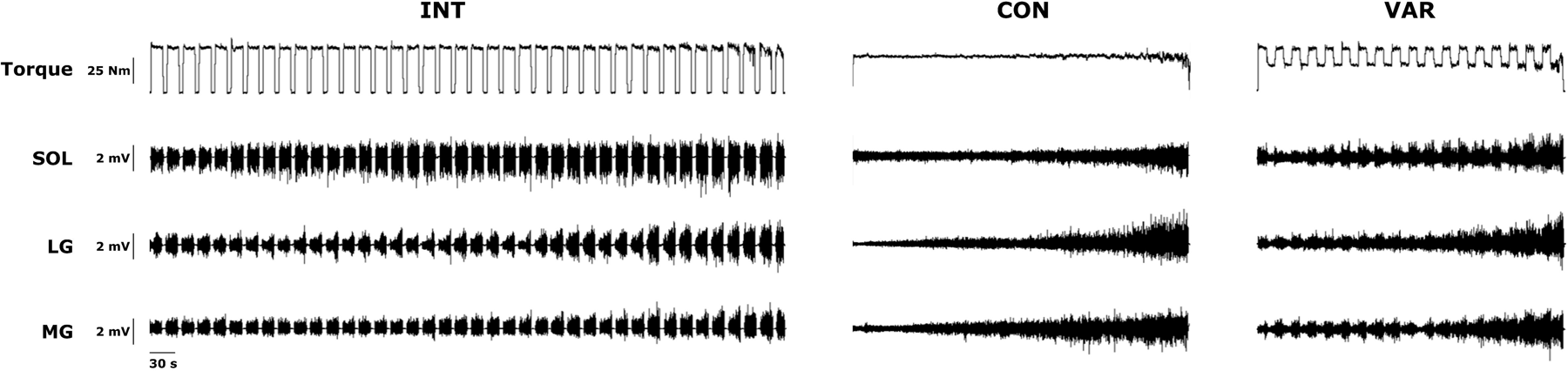
Experimental torque and electromyographic traces from a typical subject during each fatiguing exercise performed until task failure. INT = intermittent exercise (15 s on, 5 s off) at 40% MVC; CON = continuous exercise at 30% MVC; VAR = exercise with a variable contraction intensity (alternation of 10 s at 40% MVC and 10 s at 20% MVC); LG = lateral gastrocnemius; MG = medial gastrocnemius; MVC = maximal voluntary contraction; SOL = soleus

**Table 1.** Fatiguing exercise parameters.

|  | INT | CON | VAR |
| --- | --- | --- | --- |
| **Total ED (s)** | 1042.9 ± 546.0 | 506.2 ± 200.7*** | 538.4 ± 270.1*** |
| **Effective ED (s)** | 783.4 ± 408.8 | 506.2 ± 200.7*** | 538.4 ± 270.1** |
| **Exercise TTI (Nm·s)** | 35836 ± 14117 | 17492 ± 5432*** | 19187 ± 8542*** |
| **MET (Nm)** | 36.2 ± 5.7 | 35.8 ± 5.0 | 37.2 ± 6.5 |
| **MET (% MVC)** | 28.9 ± 0.8 | 28.4 ± 1.5 | 28.7 ± 1.1 |
| **RMS/M<sub>MAX</sub> beginning (x10<sup>-3</sup>)** | 9.05 ± 2.98 | 5.84 ± 1.90** | 6.45 ± 1.70* |
| **RMS/M<sub>MAX</sub> end (x10<sup>-3</sup>)** | 13.03 ± 4.36 | 12.88 ± 2.96 | 12.07 ± 3.58 |
| **RPE** | 97.8 ± 7.4 | 100.1 ± 7.2 | 95.5 ± 4.3 |

CON = continuous exercise; ED = exercise duration; INT = intermittent exercise; MET = mean exercise torque; M<sub>MAX</sub> = sum of triceps surae maximal M-wave; RMS = sum of triceps surae electromyographic root mean square; RPE = rate of perceived exertion; VAR = variable exercise. The RMS/M<sub>MAX</sub> ratio was calculated at both the beginning and the end of the fatiguing exercise. *n* = 13. \**p* < 0.05, \*\**p* < 0.01 and \*\*\**p* < 0.001 represent significant difference from INT value

At TF, the RPE did not differ among exercises (F_(2,24)_ = 3.109, *p* = 0.063, η_p_² = 0.206) and ranged between “extremely strong” and “maximal” on the CR-100 scale (Table 1).

### Assessments related to maximal torque production

*MVC*. Only a significant main *time* effect was detected on MVC (F_(2,24)_ = 145.033, *p* < 0.001, η_p_² = 0.924). Fatiguing exercises resulted in a 29.7% reduction in MVC (127.0 ± 20.7 Nm *vs*. 89.8 ± 17.7 Nm, *p* < 0.001). After the 10-min rest period, the MVC (115.4 ± 22.3 Nm) remained 10.2% below the pre-exercise value (*p* < 0.001), revealing an incomplete recovery from the fatiguing exercise (*p* < 0.001) (Figure 4A). Bayesian equivalence tests showed that MVC was likely similar among exercises at Pre (BF^OH^_01_ > 3.111 and BF^NOH^_01_ > 3.925 in all cases), Post (BF^OH^ > 2.510 and BF^NOH^_01_ > 2.947 in all cases) and Post-10 (BF^OH^_01_ > 3.226 and BF^NOH^_01_ > 4.130 in all cases).

**Figure 4.**
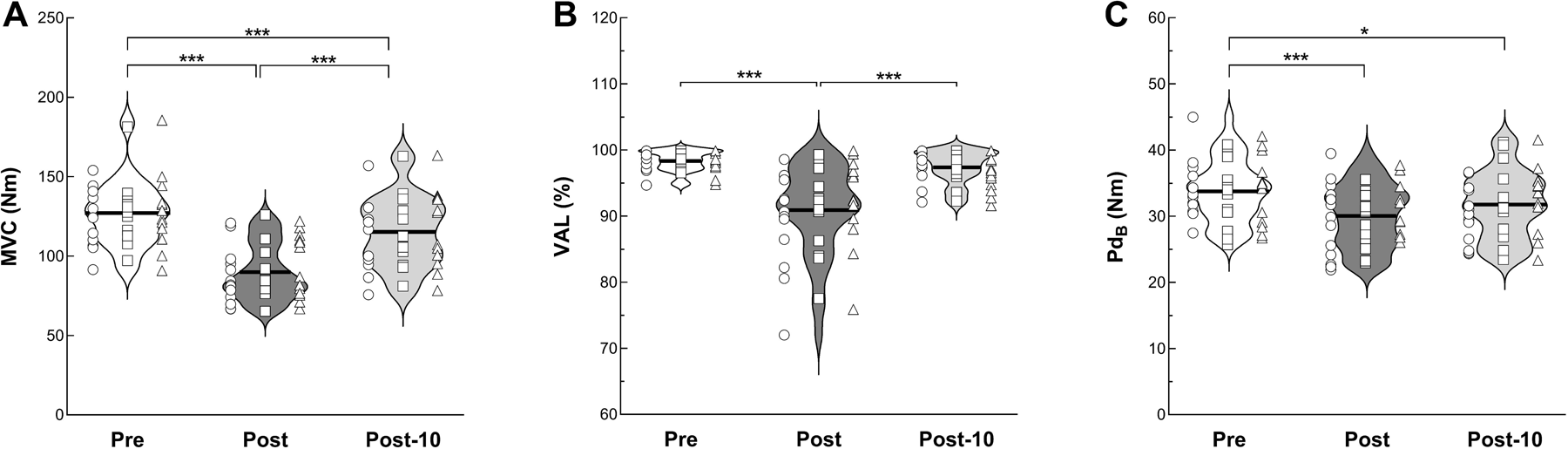
Maximal torque production, voluntary activation and potentiated doublet measured before (Pre, white violin plots) and after (Post, dark grey violin plots) the fatiguing exercise, as well as after a 10-min recovery period (Post-10, light grey violin plots). Pooled data from the three exercises are violin plotted at each time point and the horizontal black lines represent mean values. Individual data from INT (circles), CON (squares) and VAR (triangles) exercises are also presented (*n* = 13). **A**. Changes in MVC. **B**. Changes in VAL computed at the beginning of the MVC_1-MIN_. **C**. Changes in Pd_B_. CON = continuous exercise; INT = intermittent exercise; MVC = maximal voluntary contraction; MVC_1-MIN_ = sustained MVC for 1 min; Pd_B_ = potentiated doublet before the MVC_1-MIN_; VAL = voluntary activation level; VAR = variable exercise. \**p* < 0.05, \*\*\**p* < 0.001 express a significant difference

*VAL*. The ANOVA analysis only revealed a main *time* effect on VAL (F_(2,24)_ = 21.140, *p* < 0.001, η_p_² = 0.638). The VAL was decreased by 7.7% after the fatiguing exercises (98.4 ± 1.6 % *vs*. 90.8 ± 6.8 %, *p* < 0.001). The VAL fully recovered after the rest period since at Post-10, no more difference was detected with the pre-exercise value (*p* = 0.721) (Figure 4B).

*M_MAX_*. A main *time* effect was observed on the M_MAX_ amplitude (F_(2,24)_ = 18.833, *p* < 0.001, η_p_² = 0.611). The M_MAX_ following the fatiguing exercises (18.8 ± 6.9 mV) was 17.7% lower than before (22.9 ± 7.6 mV, *p* < 0.001), and fully recovered after the 10- min rest period (21.5 ± 7.4 mV) since there was no more difference with Pre-exercise (*p* = 0.102).

*Pd_B_*. Only a main *time* effect was detected on Pd_B_ (F_(2,24)_ = 12.739, *p* < 0.001, η_p_² = 0.515). The Pd_B_ decreased by 10.9% at Post (33.8 ± 4.7 Nm *vs*. 30.1 ± 4.8 Nm, *p* < 0.001). At Post-10, the Pd_B_ (31.8 ± 5.0 Nm) did not recover since it was still significantly lower than before the exercise (5.9%, *p* = 0.030) (Figure 4C).

### Assessments related to maximal torque sustainability

Δ*MVC_1-MIN_.* The ANOVA revealed only a main *time* effect on ΔMVC_1-MIN_ (F_(2,24)_ = 9.542, *p* < 0.001, η_p_² = 0.443). Compared to before (-44.9 ± 10.1 % MVC), the ΔMVC_1-MIN_ increased by 13.1% after the fatiguing exercises (-50.8 ± 10.4 % MVC, *p* = 0.001) and was fully recovered at Post-10 (-46.2 ± 9.0 % MVC) since no more difference was found with Pre-exercise (*p* = 0.617) (Figure 5A). Bayesian equivalence tests showed that ΔMVC_1-MIN_ were likely similar among exercises at Pre (BF^OH^_01_ > 3.501 and BF^NOH^_01_ > 4.641 in all cases), Post (BF^OH^_01_ > 1.711 and BF^NOH^_01_ > 1.840 in all cases) and Post-10 (BF^OH^_01_ > 1.241 and BF^NOH^_01_ > 1.271 in all cases).

**Figure 5.**
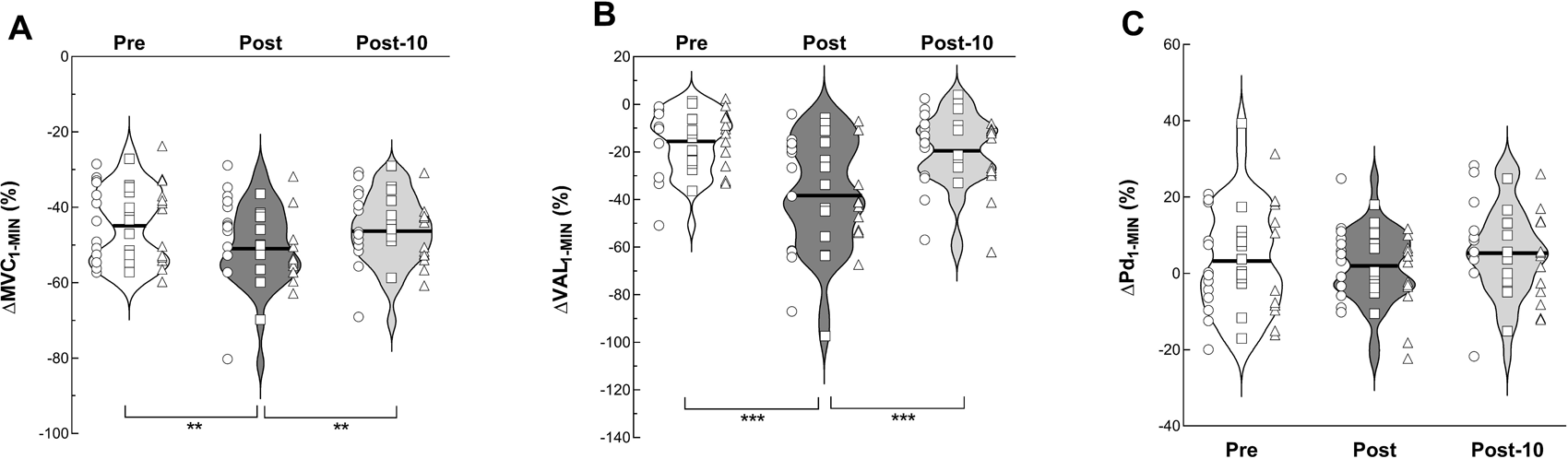
Maximal torque sustainability and associated variables measured before (Pre, white violin plots) and after (Post, dark violin plots) the fatiguing exercises, and following a 10-min recovery period (Post-10, light grey violin plots). Pooled data from the three exercises are violin plotted at each time point and the horizontal black lines represent mean values. Individual data from INT (circles), CON (squares) and VAR (triangles) exercises are also presented (*n* = 13). **A.** ΔMVC_1-MIN_ represents the percentage of torque loss during each MVC_1-MIN_. **B**. ΔVAL_1-MIN_ represents the percentage of VAL change during each MVC_1-MIN_. **C**. ΔPd_1-MIN_ represents the percentage of change in Pd during each MVC_1-MIN_. CON = continuous exercise; INT = intermittent exercise; MVC_1-MIN_ = 1-minute maximal voluntary contraction; Pd = potentiated doublet; VAL = voluntary activation level; VAR = variable exercise. \*\**p* < 0.01, \*\*\**p* < 0.001 express a significant difference

Δ*VAL_1-MIN_.* The statistical analysis showed only a main *time* effect on ΔVAL_1-MIN_ (F_(2,24)_ = 31.765, *p* < 0.001, η_p_² = 0.726). ΔVAL_1-MIN_ was 250.8% higher after the exercises (-38.2 ± 22.8 %) compared to before (-15.2 ± 12.6 %, *p* < 0.001). At Post- 10, the ΔVAL_1-MIN_ was fully recovered since no more difference was detected with the Pre value (*p* = 0.335) (Figure 5B).

Δ*Pd_1-MIN_.* No *pattern* (F_2,24_ = 0.792, *p* = 0.465, η_p_² = 0.062), *time* (F_2,24_ = 1.134, *p* = 0.339, η_p_² = 0.086) or *pattern* x *time* interaction (F_4,48_ = 1.165, *p* = 0.338, η_p_² = 0.089) effect on ΔPd_1-MIN_ was detected by statistical analysis (Figure 5C). Pre, Post and Post-10 pooled values were 3.3 ± 13.9 %, 2.0 ± 9.5 % and 5.4 ± 11.5 % respectively.

## DISCUSSION

The current study aims, using different contraction patterns, to gain a better understanding of how the mean torque of exhausting exercises affects neuromuscular fatigability. Our findings demonstrate that INT, CON and VAR contractions produce similar levels of neuromuscular fatigability, as evidenced by the comparable reductions in maximal torque production and sustainability across the exercises. These results highlight the independence of neuromuscular fatigability from the contraction pattern during such exercises and support the relationship between the MET and maximal torque sustainability. The different recovery kinetics emphasize that maximal torque production and sustainability represent distinct capacities and, therefore, must be evaluated simultaneously when exploring neuromuscular fatigue.

### Parameters characterizing fatiguing exercise

The inverse relationship between the ED and the intensity of a sustained contraction has been substantiated by previous studies (Rohmert 1960; Shahidi and Mathieu 1995; Rudroff et al. 2007). Our results allow the extension of this relationship to sustained contraction with variable intensity (i.e., VAR). Therefore, it appears that the ED of a maintained submaximal contraction correlates to MET even if the exerted torque is modulated during the exercise. This relationship is not valid anymore when integrating the INT exercise, regardless of whether we consider total or effective ED. The ED and TTI were greater for INT compared to CON and VAR. As previously reported by Lebesque et al. (2023), the presence of rest periods during exercise appears to be crucial to delay exhaustion and thus to stay in exercise longer. Indeed, rest periods during INT allow the delivery of blood flow to the active muscles, leading to a greater supply of substrates and oxygen as well as an enhanced metabolite washout (Sahlin 1992; Degens et al. 1998; Hargreaves 2005; Ament and Verkerke 2009). It is noteworthy that modulating the contraction intensity from 40 to 20% MVC during the VAR is not sufficient to extend the ED and TTI since, for both parameters, no difference was observed with CON.

Since, at the beginning of the fatiguing exercise, the torque developed during the INT (40% MVC) was greater than CON (30% MVC) and VAR (20 and 40% MVC, mean torque = 30% MVC), for this modality of exercise we also observed a greater PF electrical activity. The PF EMG activity increased from contraction onset to task failure in all exercises, denoting a rise in the excitatory output of the motor neurons (Fuglevand et al. 1993; Søgaard et al. 2006; Klass et al. 2008). This could be explained by the increased recruitment of motor units and the modulation of their firing rate to maintain the required torque, despite the development of neuromuscular fatigability (Garland et al. 1994; Carpentier et al. 2001; Kallenberg and Hermens 2006; Enoka and Duchateau 2008; Klass et al. 2008). Regardless of the contraction pattern, in our study, it appears that exhaustion was achieved for a similar level of PF EMG activity, suggesting that TF occurs when the voluntary descending drive mismatches the requirement to maintain the target torque. This finding is in accordance with the recent data from Matta et al. (2023). By manipulating the time perception during exercise, these authors highlight that subjects stopped their CON contraction of the knee extensors when a critical value of EMG activity was reached, independently of ED. Furthermore, in the present study, the effective ED negatively correlates with the change in muscle activity during exhausting exercises, showing that the greater the increase in muscle activity, the lower the ED. These findings are in agreement with those observed during submaximal CON contractions sustained until TF (Maluf et al. 2005; Hunter et al. 2008; Rudroff et al. 2011).

In the current investigation, the RPE quantified at TF was similar among exercises. This common observation at the end of exhausting exercises (e.g. Hunter et al. 2005, 2008; Rudroff et al. 2011; Williams et al. 2014) suggests that participants performed a similar subjective maximal performance during INT, CON and VAR exercises, irrespective of exercise duration or TTI.

### Maximal torque production

In the present investigation, the maximal torque production was quantified as the maximal torque achieved during the first 5 s of the MVC_1-MIN_. This methodology for assessing maximal torque production from a sustained MVC has already been used by our research group (Lebesque et al. 2022, 2023). Indeed, it has been shown that MVC from a MVC_1-MIN_ is a reliable and reproducible index of maximal torque production when compared to the brief-MVC (Lebesque et al. 2022, 2023). As hypothesized in the present study, the MVC loss after each exhausting exercises was similar (-30 %), regardless of the ED or the TTI. In one of our recent investigations, we also found a similar reduction in MVC (-27%) following CON and INT contractions performed at 40% MVC until TF (Lebesque et al. 2023). Similarly, Smith et al. (2016) observed an equivalent reduction in knee extensors MVC (-24%) after 50 intermittent contractions (6 s on, 2 s off) performed at a constant (60% MVC) or variable (i.e., alternation of contractions at 40 and 80% MVC) intensity. Altogether, these outcomes bring out that the reduction of maximal torque production is not sensitive to the contraction pattern (i.e., CON, INT and VAR).

In the present investigation, we also found a lack of correlation between the TTI and the MVC loss, for exercises performed until exhaustion with different contraction patterns. This observation could also be inferred from the results of previous investigations (Place et al. 2005, 2006a, b, 2007; Hunter et al. 2005, 2008; Klass et al. 2008; Baudry et al. 2009, 2011; Yoon et al. 2009; Rudroff et al. 2010; Neyroud et al. 2013; Williams et al. 2014), although in these latter the correlation was not specifically addressed.

The equivalent MVC decline (-30%), induced by the three fatiguing tasks in the present experiment, was associated with a reduction in VAL (-8%) which was also equivalent among exercises. This finding suggests that the central nervous system was similarly affected by the three contraction patterns and was no longer able to maximally activate muscles at TF. Prior studies, with different protocols, have noted a voluntary activation decrease following CON (e.g. Place et al. 2007; Neyroud et al. 2013) or INT (e.g. Bigland-Ritchie et al. 1986; Behm and St-Pierre 1998; Mademli and Arampatzis 2008) exercises performed until TF at 40 – 65 % MVC, but our recent investigation (Lebesque et al. 2023) stands out as the first to directly compare voluntary activation after these two modalities of contraction showing an equivalent impact (-11%) of both on VAL.

In the current research, the decline in MVC is also associated with alterations occurring at the muscular level, as revealed by the decline in Pd_B_ (-11%) and M_MAX_ (- 18%). The extent of these impairments was equivalent for the three fatiguing exercises and demonstrated that alteration in muscle contractility can mainly be explained by the decline in sarcolemma excitability.

Altogether, INT, CON and VAR contractions, performed until exhaustion, induced a similar reduction in maximal torque production that may be attributed to both neural and muscular impairments, with a larger contribution from the latter. These alterations are quantitatively similar among exercises.

The effect of a 10-min recovery period on the neuromuscular function was also investigated. We observed that the maximal torque production recovered only partially at Post-10 since the MVC remained lower, at this time point, compared to the pre-exercise value. This finding agrees with previous investigations showing that MVC was still depressed 10 min after CON (Fuglevand et al. 1993; Lebesque et al. 2023) or INT (Mademli and Arampatzis 2008; Lebesque et al. 2023) contractions performed until exhaustion. Specifically, our results suggest that the incomplete recovery of maximal torque production was mainly due to muscular factors since, regardless of the contraction pattern, the VAL fully recovered at Post-10, while the Pd_B_ was still lower than the pre-exercise value. Since the sarcolemma excitability (i.e., M_MAX_) was restored after the recovery period, it can be inferred that the alterations occurred, more precisely, within the muscle cell (Allen et al. 2008; Cheng et al. 2018). These findings suggest that intracellular mechanisms including calcic phenomena, might be involved in the alteration of muscle contractility (i.e., Pd_B_) after the fatiguing exercises (Place et al. 2007; Allen et al. 2008; Cheng et al. 2018).

### Maximal torque sustainability

According to our hypothesis, the INT, CON and VAR exercises similarly altered the ΔMVC_1-MIN_. This finding means that exhausting exercises performed with an identical MET lead to an equivalent impairment of the ability to sustain a maximal effort, regardless of the ED or the TTI. This outcome appears to support the significant correlation between the MET and the alteration in maximal torque sustainability (Lebesque et al. 2023). Indeed, although the contraction pattern was different among exercises, the MET was equivalent and it led to a similar decline in maximal torque sustainability. If such a relationship did not exist, we should have detected a different alteration in maximal torque sustainability among the exercises.

The similar alteration of maximal torque sustainability, after the three exercises, was accompanied by a similar increase in ΔVAL_1-MIN_. The accumulation of contraction by-products in the intracellular milieu is known to stimulate group III/IV muscle afferents through receptors sensitive to the mechanic and metabolic changes (Garland and Kaufman 1995), leading to the increase of motor cortex inhibition that limits the efferent motor command (Gandevia et al. 1996; Gandevia 2001; Taylor et al. 2016).

The absence of change in ΔPd_1-MIN_ stresses that the decline in maximal torque sustainability may be mainly explained by the reduction in the capacity to maintain a high level of voluntary activation, confirming our previous finding following CON and INT exhausting exercises (Lebesque et al. 2023). This observation has also be made after more ecological exercises such as cycling (Girard et al. 2013). The link between the capacities to maintain the maximal torque and the maximal voluntary activation is also hinted by the fact that the 10 min recovery period was sufficient to restore both the ΔVAL_1-MIN_ and the ΔMVC_1-MIN_.

Regardless of the contraction pattern, the neuromuscular fatigability is similar when fatiguing exercises are performed until TF. Indeed, the ability to produce and sustain the maximal torque was altered to the same extent at the end of the different exercises. However, the behaviour of MVC and ΔMVC_1-MIN_ was different after the rest of 10 min. Contrary to the MVC that did not completely recover after the rest period, the ΔMVC_1-MIN_ fully recovered at Post-10. These results are in accordance with previous observations done 10 min after isometric exercises of PF performed to exhaustion (Lebesque et al. 2023) or not (Lebesque et al. 2022). The present data therefore confirm that maximal torque production and sustainability represent distinct neuromuscular capacities. Accordingly, it is not possible to extrapolate the behaviour of one of these from the other, as experimentally shown by the absence of correlation between MVC and ΔMVC_1-MIN_ changes after the fatiguing exercises and following the recovery period (see Supplemental Figure S1). Therefore, it is important to evaluate both simultaneously when aiming to explore more deeply the neuromuscular fatigability of a muscle task or the effects of a training program on neuromuscular function.

## Conclusion

In the present investigation, we demonstrated that INT, CON and VAR submaximal exercises, performed until exhaustion with an equivalent MET, lead to a similar reduction in both maximal torque production and sustainability. Therefore, by evaluating both capacities (Lebesque et al. 2022, 2023), it can be concluded that these exercises induce the same neuromuscular fatigability. In addition, the distinct recovery kinetic of maximal torque production and sustainability confirms the importance of assessing both capacities simultaneously to get a more comprehensive description of neuromuscular fatigability. Finally, our results show a correlation between MET and change in ΔMVC_1-MIN_, namely a sensitivity of maximal torque sustainability to MET. This finding is crucial to consider for the management of neuromuscular fatigability during exercise training and reconditioning in both athletes and pathological persons.

## ACKNOWLEDGMENTS

The authors would like to thank all volunteers that participated in the study.

## ABBREVIATIONS

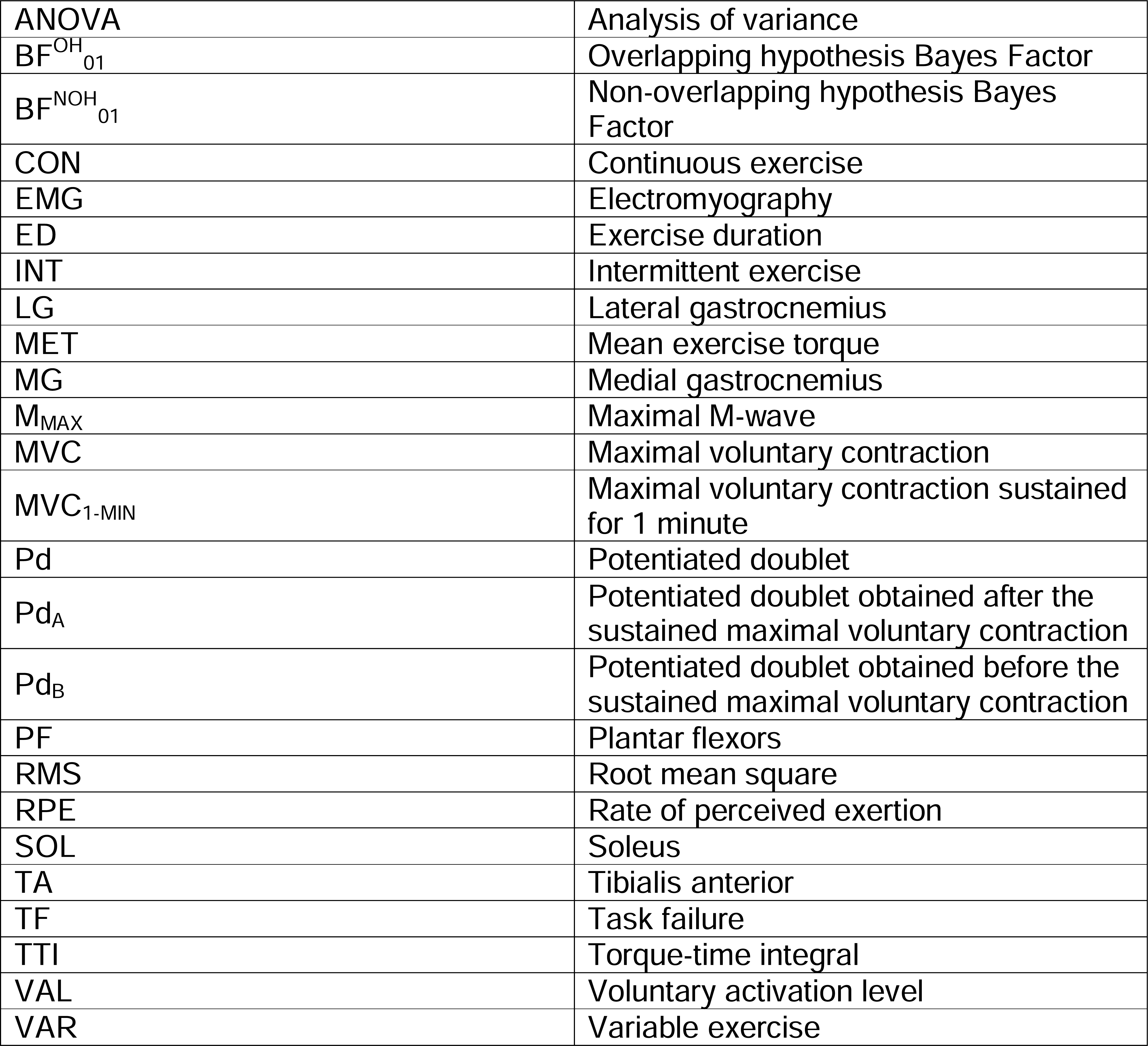

## STATEMENTS AND DECLARATIONS

### Grants

This work was supported by the Regional Council of Bourgogne Franche-Comté.

### Conflict of interest

The authors declare that the research was conducted in the absence of any commercial or financial relationship that could be construed as a potential conflict of interest.

### Data Availability

Data will be made available on reasonable request.

### Supplementary data

Supplementary data: https://doi.org/10.6084/m9.figshare.24867426

## Notes

https://doi.org/10.6084/m9.figshare.24867426

### Competing Interest Statement

The authors have declared no competing interest.

### Summary of Updates

Adding an author in the submission form; no change in the manuscript file.

