## Supplementary data for "Mean exercise torque is a critical factor influencing neuromuscular fatigability induced by exhausting contractions"

**JOURNAL NAME:** European Journal of Applied Physiology

**AUTHOR NAMES:** Loïc Lebesque^1^*, Gil Scaglioni^1^, Patrick Manckoundia^1,2^, Alain Martin^1^

**AFFILIATION:** ^1^ INSERM UMR1093-CAPS, Université de Bourgogne, UFR des Sciences du Sport, F-21000, Dijon ; ^2^ Geriatrics Internal Medicine Department, University Hospital of Dijon Bourgogne, Dijon CEDEX, 21079

**Table S1** Data from neuromuscular assessment before and after the fatiguing exercise and following the recovery period (*n* = 13)

| **Variables** | **INT** | | | **CON** | | | **VAR** | | |
| --- | --- | --- | --- | --- | --- | --- | --- | --- | --- |
|  | **Pre** | **Post** | **Post-10** | **Pre** | **Post** | **Post-10** | **Pre** | **Post** | **Post-10** |
| **MVC (Nm)** | 125.0 ± 18.4 | 86.7 ± 17.8 | 113.0 ± 21.7 | 126.5 ± 20.5 | 89.3 ± 16.3 | 117.0 ± 22.5 | 129.6 ± 24.2 | 93.5 ± 19.7 | 116.3 ± 24.3 |
| **VAL (%)** | 98.0 ± 1.8 | 89.1 ± 7.2 | 98.0 ± 2.5 | 98.7 ± 1.4 | 90.9 ± 6.4 | 97.7 ± 2.5 | 98.4 ± 1.7 | 92.5 ± 6.8 | 96.4 ± 2.6 |
| **M_MAX_ (mV)** | 25.08 ± 7.50 | 21.55 ± 7.71 | 23.36 ± 8.19 | 22.53 ± 7.65 | 18.03 ± 7.61 | 21.29 ± 7.56 | 21.08 ± 7.64 | 16.93 ± 4.69 | 19.74 ± 6.44 |
| **Pd_B_ (Nm)** | 34.1 ± 4.4 | 29.5 ± 4.7 | 30.3 ± 5.0 | 33.4 ± 4.9 | 29.5 ± 4.2 | 32.2 ± 5.7 | 33.9 ± 5.2 | 31.5 ± 4.3 | 32.9 ± 4.9 |
| **∆MVC_1-MIN_ (% MVC)** | -44.7 ± 10.4 | -47.2 ± 12.8 | -45.9 ± 10.7 | -45.1 ± 9.4 | -52.1 ± 9.4 | -43.8 ± 7.7 | -45.0 ± 11.2 | -53.0 ± 8.4 | -49.0 ± 8.1 |
| **∆VAL_1-MIN_ (% VAL)** | -6.7± 8.4 | -11.7 ± 8.5 | -8.7 ± 8.4 | -8.2 ± 6.1 | -11.3 ± 8.3 | -8.5 ± 7.4 | -7.6 ± 7.0 | -14.9 ± 11.0 | -10.2 ± 7.1 |
| **∆Pd_1-MIN_ (% Pd_B_)** | 1.6 ± 12.8 | 3.9 ± 9.8 | 8.5 ± 12.5 | 5.0 ± 13.9 | 3.4 ± 8.5 | 3.8 ± 10.6 | 3.3 ± 15.8 | -1.2 ± 10.1 | 3.8 ± 11.7 |

Data are expressed as mean ± standard deviation. CON = continuous exercise; INT = intermittent exercise; VAR = variable exercise; Pre = before the fatiguing exercise; Post = after the fatiguing exercise; Post-10 = after a 10-min rest period following the fatiguing exercise; MVC = maximal voluntary contraction; ΔMVC_1-MIN_ = torque loss during the 1-min sustained maximal voluntary contraction; VAL = voluntary activation level; ΔVAL_1-MIN_ = change in voluntary activation level between the beginning and the end of the sustained maximal voluntary contraction; Pd = potentiated doublet; ΔPd_1-MIN_ = change in potentiated doublet between before and after the sustained maximal voluntary contraction; M_MAX_ = sum of maximal M-wave amplitude of soleus and gastrocnemii muscles.

**Table S2** Statistic outcomes from neuromuscular variables (*n* = 13)

| **Variables** | **Normal distribution** | **Statistic test** | **Statistic outcomes** |
| --- | --- | --- | --- |
| **MVC (Nm)** | Yes | rmANOVA (*Pattern*, *Time*)  Bayesian equivalence test | *Pattern* : F_(2,24)_ = 1.184, *p* = 0.323, *η*_p_² = 0.090    *Time* : F_(2,24)_ = 145.033, *p* < 0.001, *η*_p_² = 0.924  *Pattern* x *Time* : F_(4,48)_ = 0.516, *p* = 0.724, *η*_p_² = 0.041  Pre : BF^OH^_01_ > 3.111, BF^NOH^_01_ > 3.928  Post : BF^OH^_01_ > 2.510, BF^NOH^_01_ > 2.947  Post-10 : BF^OH^_01_ > 3.226, BF^NOH^_01_ > 4.130 |
| **VAL (%)** | Yes | rmANOVA (*Pattern*, *Time*) | *Pattern* : F_(2,24)_ = 0.490, *p* = 0.616, *η*_p_² = 0.040    *Time* : F_(2,24)_ = 21.140, *p* < 0.001, *η*_p_² = 0.638  *Pattern* x *Time* : F_(4,48)_ = 2.310, *p* = 0.072, *η*_p_² = 0.161 |
| **M_MAX_ (mV)** | Yes | rmANOVA (*Pattern*, *Time*) | *Pattern* : F_(2,24)_ = 2.952, *p* = 0.071, *η*_p_² = 0.197    *Time* : F_(2,24)_ = 18.833, *p* < 0.001, *η*_p_² = 0.611  *Pattern* x *Time* : F_(4,48)_ = 1.089,  *p* = 0.372, *η*_p_² = 0.083 |
| **Pd_B_ (Nm)** | Yes | rmANOVA (*Pattern*, *Time*) | *Pattern* : F_(2,24)_ = 1.247, *p* = 0.305, *η*_p_² = 0.094    *Time* : F_(2,24)_ = 12.739, *p* < 0.001, *η*_p_² = 0.515  *Pattern* x *Time* : F_(4,48)_ = 2.252,  *p* = 0.077, *η*_p_² = 0.158 |
| **∆MVC_1-MIN_ (% MVC)** | Yes | rmANOVA (*Pattern*, *Time*)  Bayesian equivalence test | *Pattern* : F_(2,24)_ = 1.564, *p* = 0.230, *η*_p_² = 0.115    *Time* : F_(2,24)_ = 9.542, *p* < 0.001, *η*_p_² = 0.443  *Pattern* x *Time* : F_(4,48)_ = 1.629,  *p* = 0.182, *η*_p_² = 0.120  Pre : BF^OH^_01_ > 3.501, BF^NOH^_01_ > 4.641  Post : BF^OH^_01_ > 1.711, BF^NOH^_01_ > 1.840  Post-10 : BF^OH^_01_ > 1.241, BF^NOH^_01_ > 1.271 |
| **∆VAL_1-MIN_ (% VAL)** | Yes | rmANOVA (*Pattern*, *Time*) | *Pattern* : F_(2,24)_ = 1.105, *p* = 0.348, *η*_p_² = 0.084    *Time* : F_(2,24)_ = 31.765, *p* < 0.001, *η*_p_² = 0726  *Pattern* x *Time* : F_(4,48)_ = 0.460,  *p* = 0.765, *η*_p_² = 0.037 |
| **∆Pd_1-MIN_ (% Pd_B_)** | Yes | rmANOVA (*Pattern*, *Time*) | *Pattern* : F_(2,24)_ = 0.792, *p* = 0.465, *η*_p_² = 0.062    *Time* : F_(2,24)_ = 1.134, *p* = 0.339,  *η*_p_² = 0.086  *Pattern* x *Time* : F_(4,48)_ = 1.165,  *p* = 0.338, *η*_p_² = 0.089 |

MVC = maximal voluntary contraction; ΔMVC_1-MIN_ = torque loss during the 1-min sustained maximal voluntary contraction; VAL = voluntary activation level; ΔVAL_1-MIN_ = change in voluntary activation level between the beginning and the end of the sustained maximal voluntary contraction; Pd = potentiated doublet; ΔPd_1-MIN_ = change in potentiated doublet between before and after the sustained maximal voluntary contraction; M_MAX_ = sum of maximal M-wave amplitude of soleus and gastrocnemii muscles; rmANOVA = repeated-measures analysis of variance with *Pattern* (INT *vs.* CON *vs.* VAR) and *Time* (Pre *vs.* Post *vs.* Post-10) factors; BF^OH^_01_ = Overlapping hypothesis Bayes Factor; BF^NOH^_01_ = Non-overlapping hypothesis Bayes Factor. The significance level was set at *p* < 0.050.

**Table S3** Pooled data from neuromuscular assessment before and after the fatiguing exercise and following the recovery period (*n* = 13)

| **Variables** | **Pooled data (INT, CON and VAR)** | | |
| --- | --- | --- | --- |
|  | **Pre** | **Post** | **Post-10** |
| **MVC (Nm)** | 127.0 ± 20.7 | 89.8 ± 17.7 | 115.4 ± 22.3 |
| **VAL (%)** | 98.4 ± 1.6 | 90.8 ± 6.8 | 97.4 ± 2.5 |
| **M_MAX_ (mV)** | 22.90 ± 7.58 | 18.84 ± 6.93 | 21.46 ± 7.39 |
| **Pd_B_ (Nm)** | 33.8 ± 4.7 | 30.1 ± 4.8 | 31.8 ± 5.0 |
| **∆MVC_1-MIN_ (% MVC)** | -44.9 ± 10.1 | -50.8 ± 10.4 | -46.2 ± 9.0 |
| **∆VAL_1-MIN_ (% VAL)** | -7.5± 7.0 | -12.7 ± 9.2 | -9.1 ± 7.5 |
| **∆Pd_1-MIN_ (% Pd_B_)** | 3.3 ± 13.9 | 2.0 ± 9.5 | 5.4 ± 11.5 |

Since there is no effect or only a significant *Time* effect on the neuromuscular variables (see Table S2), pooled data from the three exercises are presented. Data are expressed as mean ± standard deviation. CON = continuous exercise; INT = intermittent exercise; VAR = variable exercise; Pre = before the fatiguing exercise; Post = after the fatiguing exercise; Post-10 = after a 10-min rest period following the fatiguing exercise; MVC = maximal voluntary contraction; ΔMVC_1-MIN_ = torque loss during the 1-min sustained maximal voluntary contraction; VAL = voluntary activation level; ΔVAL_1-MIN_ = change in voluntary activation level between the beginning and the end of the sustained maximal voluntary contraction; Pd = potentiated doublet; ΔPd_1-MIN_ = change in potentiated doublet between before and after the sustained maximal voluntary contraction; M_MAX_ = sum of maximal M-wave amplitude of soleus and gastrocnemii muscles.

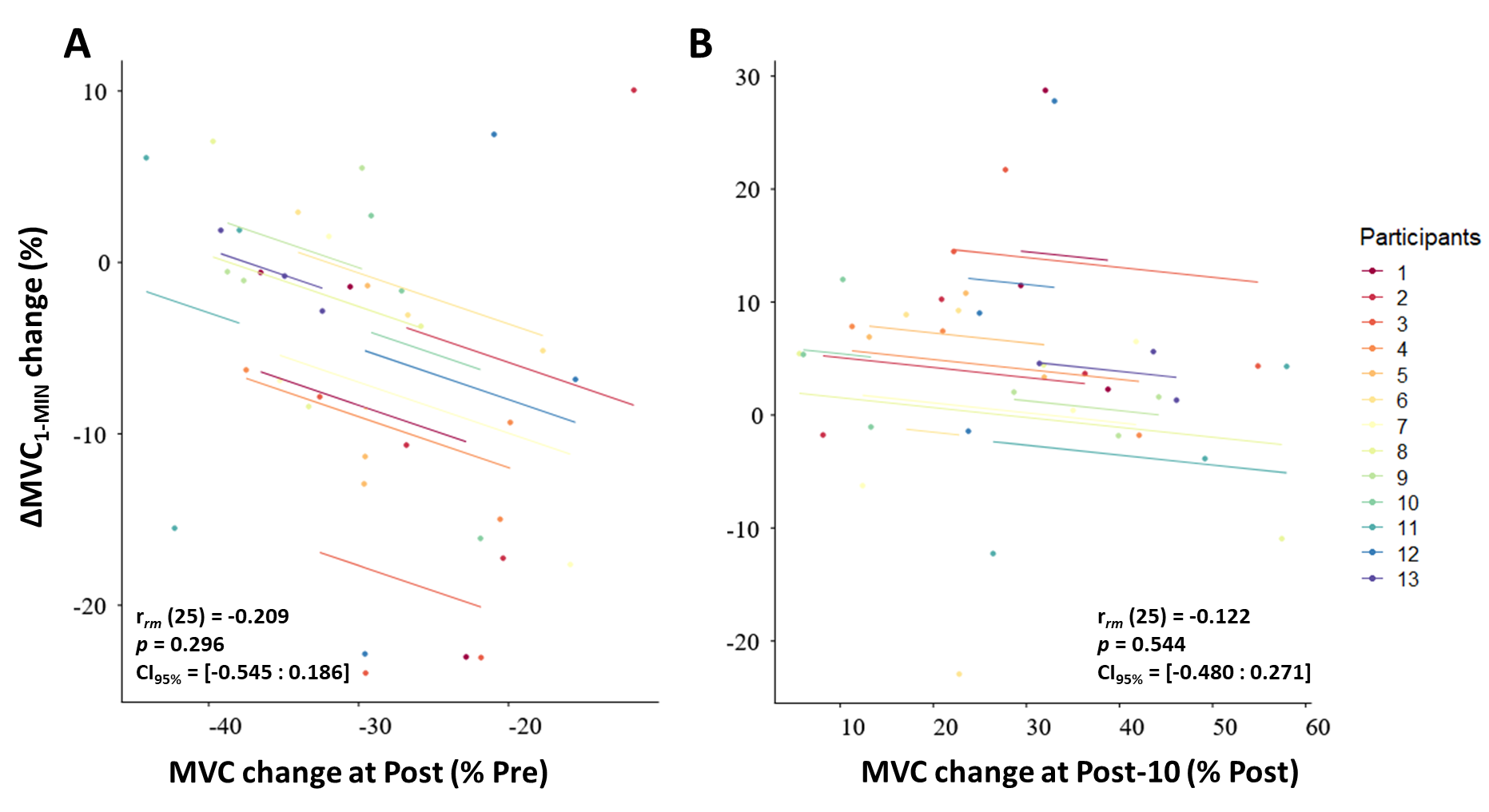

**Figure S1** Individual repeated-measures correlation between MVC and ∆MVC_1-MIN_ changes between (A) Pre and Post, and between (B) Post and Post-10. The lines correspond to the repeated-measures correlation fit for each participant (*n* = 13). For each fatiguing exercise, data from the same participant are shown in the same colour. MVC = maximal voluntary contraction; ∆MVC_1-MIN_ correspond to the relative loss of torque during a MVC sustained for 1 minute. CI_95%_ = 95% confidence interval; MVC = maximal voluntary contraction. *p* > 0.050 express a non-significant correlation.
